# Comparison of Neutralizing Antibody Titers Elicited by mRNA and Adenoviral Vector Vaccine against SARS-CoV-2 Variants

**DOI:** 10.1101/2021.07.19.452771

**Authors:** Takuya Tada, Hao Zhou, Marie I. Samanovic, Belinda M. Dcosta, Amber Cornelius, Mark J. Mulligan, Nathaniel R. Landau

## Abstract

The increasing prevalence of SARS-CoV-2 variants has raised concerns regarding possible decreases in vaccine efficacy. Here, neutralizing antibody titers elicited by mRNA-based and an adenoviral vector-based vaccine against variant pseudotyped viruses were compared. BNT162b2 and mRNA-1273-elicited antibodies showed modest neutralization resistance against Beta, Delta, Delta plus and Lambda variants whereas Ad26.COV2.S-elicited antibodies from a significant fraction of vaccinated individuals were of low neutralizing titer (IC_50_ <50). The data underscore the importance of surveillance for breakthrough infections that result in severe COVID-19 and suggest the benefit of a second immunization following Ad26.COV2.S to increase protection against the variants.

---

Severe acute respiratory syndrome coronavirus 2 (SARS-CoV-2) vaccines from two vaccine platforms have been granted U.S. Food and Drug Administration (FDA) Emergency Use Authorization: mRNA-based (Pfizer and Moderna) and adenoviral vector-based (Johnson & Johnson (J&J)), all of which have been shown to be highly effective. The mRNA-based vaccines were 94-95% effective in preventing COVID-19^1^ whereas the adenoviral vector-based J&J vaccine had 66.9% efficacy in preventing moderate to severe disease^2^. However, the ongoing emergence of highly transmissible variants with mutations in the spike protein raises concerns regarding possible decreases in vaccine effectiveness due to spike protein antigenic variability.

SARS-CoV-2 variants have been classified by the World Health Organization (WHO) based on increased transmissibility and/or pathogenicity as variants of concern (VOC; Alpha (B.1.1.7), Beta (B.1.351), Gamma (B.1.1.248) and Delta (B.1.617.2) and variants of interest (VOI; Epsilon (B.1.427/B.1.429), Iota (B.1.526), and Delta plus (AY.1) and Lambda (C.37)^3^. The increased transmissibility and/or pathogenicity of the variants is due, at least in part, to mutations in the spike protein RBD that increase its affinity for ACE2 on target cells. Mutations in the Beta, Gamma and Delta variant spike RBDs have been shown to cause partial resistance to neutralization by the serum antibodies of vaccinated and convalescent individuals and therapeutic monoclonal antibodies^4-11^.

This study compared the neutralization titers of serum antibodies from individuals immunized with three U.S. FDA Emergency use authorization vaccines (BNT162b2, mRNA-1273 and Ad26.COV2.S) against viruses with the VOC and Lambda spike proteins. The study groups were controlled for age, clinical co-morbidity, history of pre-vaccination infection and sera were collected on similar days post-vaccination. The results demonstrate a high level of cross-neutralization by antibodies elicited by BNT162b2 and mRNA-1273 on the variants but significantly decreased neutralization by those elicited by the single dose Ad26.COV2.S.

## Variant pseudotyped lentiviruses

The Delta plus spike contains K417N, L452R and T478K in the RBD **(Figure S1A)**. The Lambda spike protein contains novel L452Q and F490S mutations in the RBD **(Figure S1A)**. We previously described the production of lentiviruses pseudotyped by the Alpha, Beta, Gamma and Delta spike proteins and here report the generation of pseudotypes with the Delta plus and Lambda variant spike proteins and the individual constituent mutations. The variant spike proteins were well expressed, proteolytically processed and incorporated into lentiviral virions at a level similar to that of the parental D614G spike protein in the producer cells and virions **(Figure S1B)**. The measurement of neutralizing antibody titers with such pseudotypes has been shown to yield results consistent with those obtained with the live virus plaque reduction neutralization test^12^.

## Reduced sensitivity of virus with variant spikes to neutralization by convalescent sera and mRNA vaccine-elicited antibodies

Sera from individuals who had been infected prior to the emergence of the variants (collected 32-57 days post symptom onset) neutralized virus with the D614G spike protein with an average IC_50_ titer of 346 and neutralized the Alpha variant with a similar titer (IC_50_ of 305). Neutralizing titers for Beta, Delta, Delta plus and Lambda variants were decreased 3.2-4.9-fold relative to D614G, indicative of a modest resistance to neutralization **(Figure 1A, Table S1)**. The sera of individuals vaccinated with BNT162b2 and mRNA-1273 that were collected 7-days post-second injection – a peak antibody response timepoint-neutralized virus with the D614G spike with significantly higher titer (1835 and 1594, respectively) relative to the convalescent sera, and the antibodies cross-reacted on the variants with a modest 2.5-4.0-fold decrease in titer **(Figure 1A)**. The resistance of the Beta variant was attributed to the E484K mutation whereas resistance of the Delta variant was attributed to the L452R mutation **(Figure S2)**. The resistance of the lambda variant was attributed to both the L452Q and F490S mutations **(Figure S2)**.

**Figure 1.**
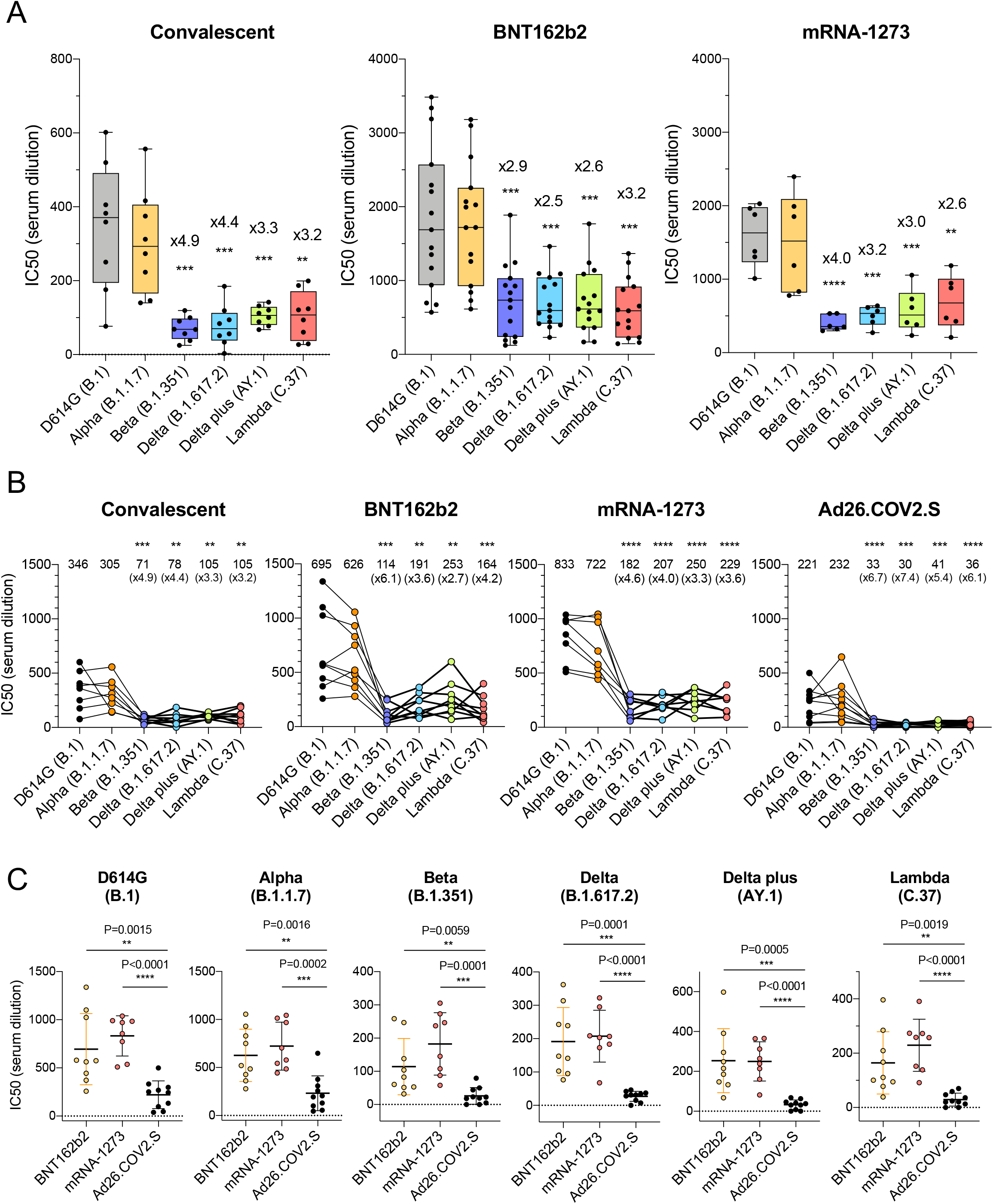
Comparison of neutralization titers of variant spike protein pseudotyped viruses by convalescent sera, antibodies elicited by BNT162b2, mRNA-1273, Ad26.COV2.S. (A) Neutralization of variant spike protein pseudotyped viruses by convalescent serum (n=8) (left). Neutralizing titers of serum samples from BNT162b2 vaccinated individuals (n=15) (middle). Neutralizing titers of serum samples from mRNA-1273 vaccinated donors (n=6) (right). The serum was collected at early time point (7 days after second immunization). The neutralization IC_50_ from individual donors is shown. Significance is based on two-sided t-test. (B) Comparison of neutralization of variants by convalescent serum (n=8, the same donors in A), BNT162b2 vaccinated individuals (n=9), mRNA-1273 vaccinated donors (n=8), Ad26.COV2.S vaccinated donors (n=10), sera from vaccinated individuals were collected at later time points (90, 80, 82 days on average after last immunization of each vaccine, see the table S2). Each line shows individual donors. (C) Comparison of neutralization potency of each vaccine by different SARS-CoV-2 variants. The neutralization IC_50_ from individual donors vaccinated by BNT162b2 (yellow), mRNA-1273 (pink), Ad26.COV2.S (black) is shown. Significance is based on two-sided t-test.

## Resistance of viruses with variant spike proteins to neutralization by Ad26.COV2.S-elicited antibodies

We next compared the neutralizing titers of antibodies elicited by the BNT162b2 and mRNA-1273 mRNA vaccines with that of the Ad26.COV2.S adenoviral vector-based vaccine. The sera analyzed were collected from individuals at similar time-points post-final injection, on average (90 days for BNT162b2, 80 days for mRNA-1273 and 82 days for Ad26.COV2.S; **Table S2**) and from individuals of similar age and with similar clinical co-morbidities **(Table S2)**. None of the participants had a history of COVID-19 pre-or post-vaccination and all were negative for antibodies against the SARS-CoV-2 N protein **(Table S2)**. The results showed that BNT162b2 sera neutralized virus with the D614G and Alpha spikes with an average titer of 695 and 626. Compared to the D614G, the neutralizing titer against Beta was decreased 6.1-fold and Delta plus was decreased 2.7-fold. Results for the mRNA-1273 vaccine were similar with a 3.3-fold decrease in neutralizing titer for Delta plus and 4.6-fold for Beta. Ad26.COV2.S sera neutralized D614G and Alpha variants with average IC_50_ titers of 221 and 232, respectively, and neutralized the variants with titers that were decreased by 5.4-fold for Delta plus to 6.7-fold for the Beta variant as compared to D614G **(Figure 1B)**. Presentation of the data grouped by variant shows the decreased neutralizing titers against the variants by sera of the Ad26.COV2.S-vaccinated individuals **(Figure 1C)**.

## The L452R/Q mutation of the Delta plus and Lambda spike proteins increases infectivity and affinity for ACE2

Measurement of the infectivity of the pseudotyped viruses, normalized for particle number, showed that the Lambda variant spike protein increased viral infectivity by 2-fold **(Figure 2A)**, an increase equivalent to that of the Delta and Delta plus variants. The increase was due to the L452Q mutation and was similar to that of the L452R found in the Delta and Delta plus variants. The other mutations (Δ246-252, G75V-T76I, F490S and T859N) had no significant effect on infectivity **(Figure 2A)**. Measurement of the relative affinity of the variant spike proteins for ACE2 using sACE2 neutralization assay showed that variant spikes had a 3-fold increase in sACE2 binding **(Figure 2B)**. This increase was confirmed in a virion:ACE2 binding assay **(Figure 2C)**. The increase was caused by the L452R and L452Q mutation and were similar to the increase caused by the N501Y mutation^13,14^.

**Figure 2.**
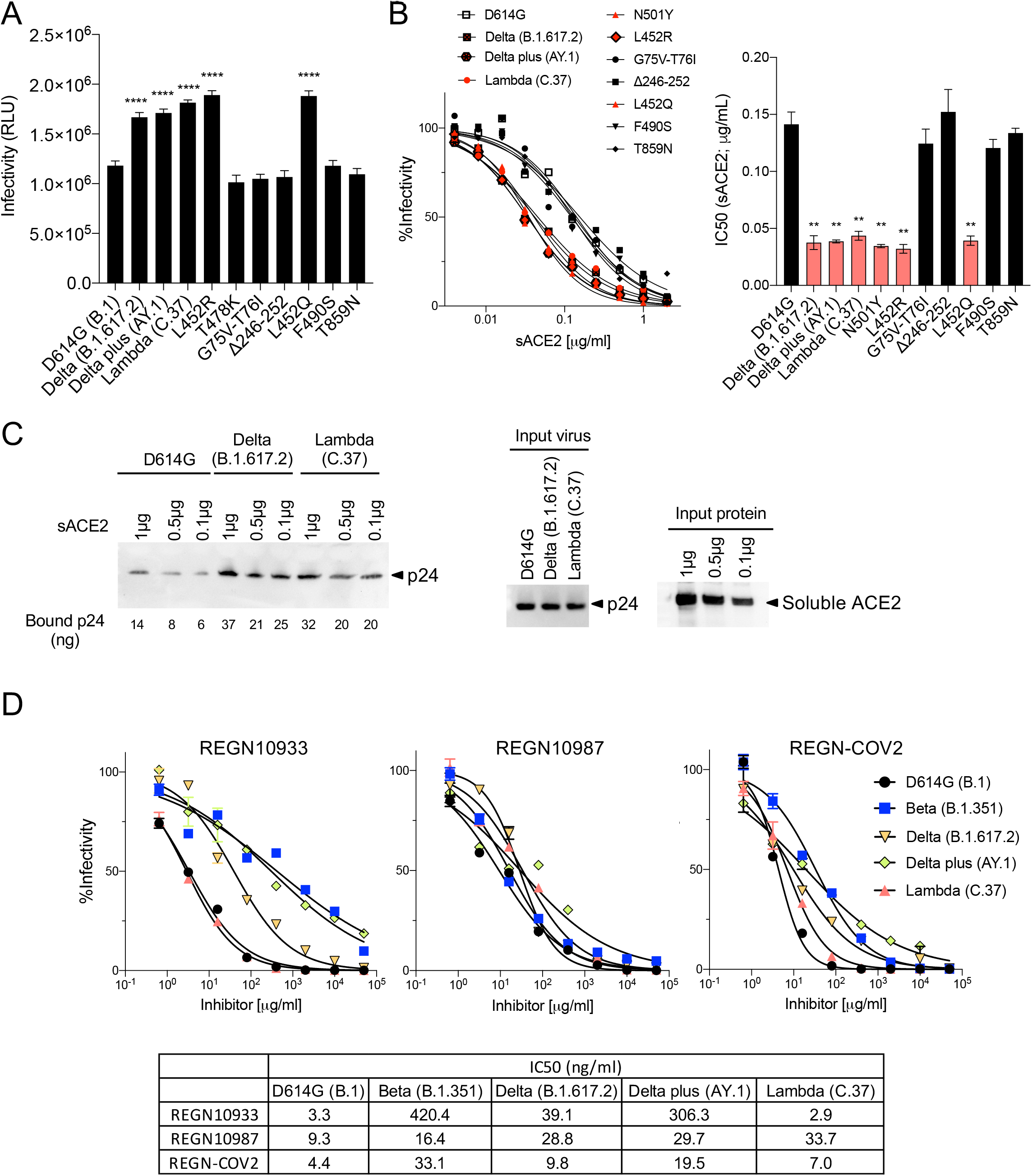
Neutralization of variant spike protein pseudotyped viruses by monoclonal antibodies and sACE2. (A) Infectivity of virus pseudotyped by variant and D614G spike proteins. Viruses were normalized for RT activity and applied to target cells. Infectivity of viruses pseudotyped with the variant proteins or the individual Lambda mutations were tested on ACE2.293T. Luciferase activity was measured two days post-infection. Significance was based on two-sided t-test. (B) Neutralization of variant spike protein variants by sACE2. Viruses pseudotyped with variant spike proteins were incubated with a serially diluted recombinant sACE2 and then applied to ACE2.293T cells. Each plot represents the percent infectivity of D614G and other mutated spike pseudotyped virus. The diagram shows the IC_50_ for each curve. (C) Nickel beads were coated for 1 hour with 1, 0.5 and 0.1 μg of sACE2 proteins. Unbound protein was removed and SARS-CoV-2 variant pseudotyped virions (D614G, Delta, Lambda) were incubated with the beads. After 1 hour, the bound virions were analyzed on an immunoblot with antibody p24 antibody. Beads-bound p24 (ng) was calculated and indicated in the bottom (left). Input virions were analyzed on an immunoblot with anti-p24 antibody (middle). Input sACE2 proteins were analyzed on an immunoblot with anti-His-tag antibody (right). (D) Neutralization of Beta, Delta, Delta plus and Lambda variant spike protein variants by REGN10933 and REGN10987 monoclonal antibodies. Neutralization of D614G and variant pseudotyped viruses by REGN10933 (left), REGN10987 (middle), and 1:1 ratio of REGN10933 and REGN10987 (right). The IC_50_ values of REGN10933, REGN10987 and the cocktail is shown in the table.

## Neutralization by REGN10933 and REGN10987

Analysis of REGN10933 and REGN10987 monoclonal antibodies that constitute the REGN-COV2 therapy showed that REGN10933 had decreased activity against the Beta variant spike which resulted in a 127-fold decrease in neutralizing titer. REGN10933 also had decreased activity against the Delta plus variant which resulted in a 92.7-fold decrease in neutralizing titer. The resistance to REGN10933 was attributed to K417N and E484K **(Figure S3)**. REGN10933 neutralized virus with the Delta variant spike with a 12-fold decrease in titer which had only a minor effect on the activity of the cocktail. REGN10987 showed a minor reduction in neutralizing titer of virus with the Beta, Delta, Delta plus and Lambda variant spikes but this had little effect on neutralization of the virus by the cocktail **(Figure 2D)**. The resistance of variants to REGN10987 was attributed to the L452R/Q **(Figure S3)**.

## Discussion

Several reports have shown partial resistance of SARS-CoV-2 VOCs to vaccine-elicited antibodies^4-11^. The data shown here extend those findings to the Delta plus and Lambda variants. Delta plus and Lambda, VOIs, both displayed a degree of resistance to mRNA vaccine-elicited antibodies similar to that of the Beta and Delta variants. In sera collected ∼3 months post-second immunization, BNT162b2 and mRNA-1273 mRNA vaccine-elicited antibodies neutralized the variants with a modest 3-fold average decrease in titer resulting in an average IC_50_ of about 1:600, a titer that is greater than that of convalescent sera and likely, in combination with post-vaccination T-and B-cell memory responses, to provide durable protection. Ad26.COV2.S vaccination-elicited neutralizing antibodies showed a more pronounced decrease in neutralizing titer against the variants, raising the potential for decreased protection against the VOCs and the Lambda variant. Vaccination with Ad26.COV2.S resulted in IC_50_ titers against Beta, Delta, Delta plus and Lambda variants that decreased 5-7-fold, resulting in mean neutralizing antibody titers of 33, 30, 41, and 36 against viruses with the Beta, Delta, Delta plus and Lambda variant spikes, respectively, which according to mathematical modeling, could result in decreased protection against infection^15^. Modeling predicts that 50% protection from infection is provided by a titer that is 20% that of the mean convalescent titer. In this study, given a mean convalescent titer of 346 **(Table S1)**, 50% protection would correspond to an IC_50_ of 69. The titer required to protect against severe disease was shown to be 3% that of the mean titer of convalescent sera which in this study corresponds to a titer of 10. In a published report of phase 3 trial data, a single dose of Ad26.COV2.S, 28 days post administration, provided 64.0% protection against moderate to severe disease and 81.7% against severe-critical COVID-19 in a country where 95% of circulating SARS-CoV-2 was the Beta variant^2^. The authors considered possible roles for non-neutralizing antibody Fc-mediated effector functions and the role of the T cell response in maintaining protection against the partially neutralizing antibody-resistant Beta variant.

The data reported here differ somewhat from those reported by Barouch *et al*. and Jongeneelen *et al*. who found that Ad26.COV2.S-elicited antibody titers were mostly maintained against the variants^16,17^. In addition, Alter *et al*. reported a 5-fold decrease in neutralizing antibody titer against Beta and 3.3-fold decrease against the Gamma variant by the sera from Ad26.COV2.S vaccination^18^ which were less pronounced than those reported here. While the studies used similar assays to measure antibody neutralization and analyzed sera collected at a similar time-point post-immunization, it is possible that differences in the study populations accounted for the experimental differences.

Several recent studies have shown that boosting a single immunization of the ChAdOx1nCoV-19 adenoviral vector vaccine with BNT162b2 resulted in high neutralizing titer against the VOCs^19-21^. It is likely that neutralizing antibody titers against the VOCs elicited by the single shot Ad26.COV2.S could similarly be improved by boosting with a second immunization or by a heterologous boost with one of the mRNA vaccines. While a single dose vaccination has advantages, the benefit provided by a second immunization may be well worth the inconvenience.

The data presented here emphasize the importance of surveillance for breakthrough infections with the increased prevalence of highly transmissible variants. If an increase in breakthrough infections accompanied by severe COVID-19 is found following adenovirus vector or mRNA vaccination, this would provide a rationale for public health policy-makers and manufacturers to consider booster immunizations that would increase protection against the VOCs and Lambda variant. As such a need is not currently evident, the public health apparatus should focus on primary immunization in the U.S. and globally.

## Methods

### Clinical Samples

Convalescent sera were collected 32-57 days post-symptom onset. For the early time-point, BNT162b2 and Moderna-vaccinated sera were collected on day 28 and 35, respectively, 7 days post-second immunization. For the later time-point, BNT162b2-vaccinated sera were on average collected 90 days post-second immunization and mRNA-1273-vaccinated sera were collected on average 80 days post-second immunization. Ad26.COV2.S-vaccinated sera were collected, on average, 82 days post-immunization **(Table S2)**. Blood was drawn at the NYU Vaccine Center with written consent under IRB approved protocols (IRB 18-02035 and IRB 18-02037). REGN10933 and REGN10987 were generated as previously described^22^.

### SARS-CoV-2 spike lentiviral pseudotypes

Lentiviruses pseudotyped by variant SARS-CoV-2 spikes were produced as previously reported^23^ and normalized for reverse transcriptase (RT) activity. Neutralization titers of sera, monoclonal antibody and soluble ACE2 (sACE2)^24^ were determined as previously described^23^.

### sACE2 pull-down assay

sACE2-bound-beads were mixed with pseudotyped virions as previously described^24^. The amount of virus bound was quantified by immunoblot analysis of bound p24.

### Statistical Analysis

All experiments were in technical duplicates or triplicates. Statistical significance was determined by two-tailed, unpaired t-test with confidence intervals shown as the mean ± SD or SEM. (*P≤0.05, **P≤0.01, ***P≤0.001, ****P≤0.0001). Spike protein structure (7BNM)^25^ was downloaded from the Protein Data Bank.

## Acknowledgements

The work was funded in part by grants from the NIH to N.R.L. (DA046100, AI122390 and AI120898) and to M.J.M. (UM1AI148574). T.T. was supported by the Vilcek/Goldfarb Fellowship Endowment Fund. M.J.M. and M.I.S. were partially supported by NYU Grossman SOM institutional support.

## Author contributions

T.T. and N.R.L. designed the experiments. H.Z., T.T. and B.M.D. carried out the experiments and analyzed data. T.T., H.Z. and N.R.L. wrote the manuscript. M.I.S. and M.J.M. designed and supervised the specimen selection, clinical information collection and the N ELISAs, and provided key reagents and useful insights. All authors provided critical comments on manuscript.

## Figure legends

**Supplemental Figure S1.**
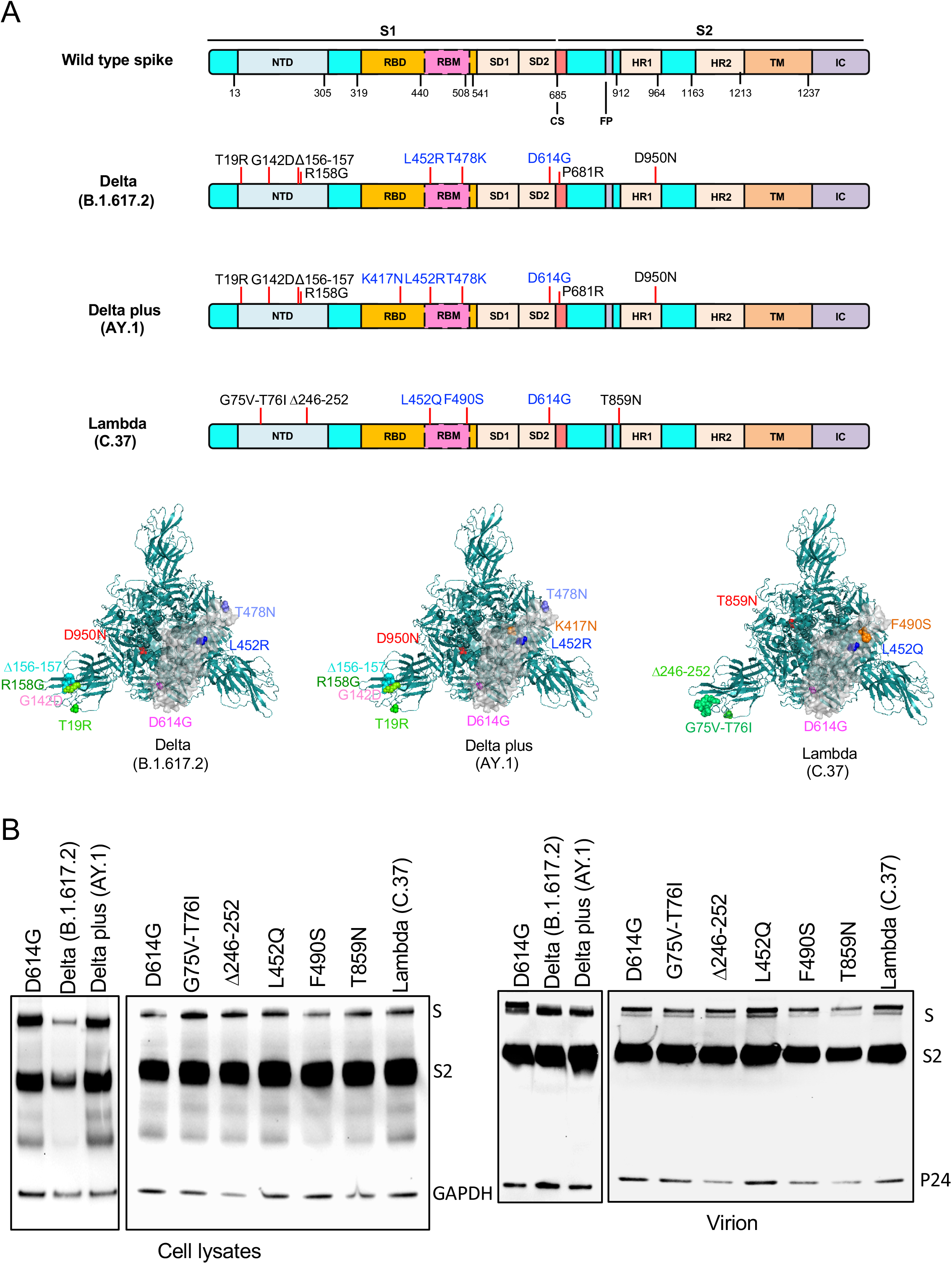
The structure of variant spikes and immunoblot analysis of spike proteins. (A) The domain structure of the SARS-CoV-2 spike is diagrammed with Delta (B.1.617.2), Delta plus (AY.1), Lambda (C.37) variant amino acid residues indicated. NTD, N-terminal domain; RBD, receptor-binding domain; RBM, receptor-binding motif; SD1 subdomain 1; SD2, subdomain 2; CS, cleavage site; FP, fusion peptide; HR1, heptad repeat 1; HR2, heptad repeat 2; TM, transmembrane region; IC, intracellular domain. Key mutations are shown in 3D structure (top view). (B) Immunoblot analysis of the Delta (B.1.617.2), Delta plus (AY.1), single point mutated of Lambda (C.37) variant, Lambda (C.37) variant spike proteins in transfected 293T cells. Pseudotyped viruses were produced by transfection of 293T cells. Two days post-transfection, virions were analyzed on an immunoblot probed with anti-spike antibody and anti-HIV-1 p24. The cell lysates were probed with anti-spike antibody and anti-GAPDH antibodies as a loading control.

**Supplemental Figure S2.**
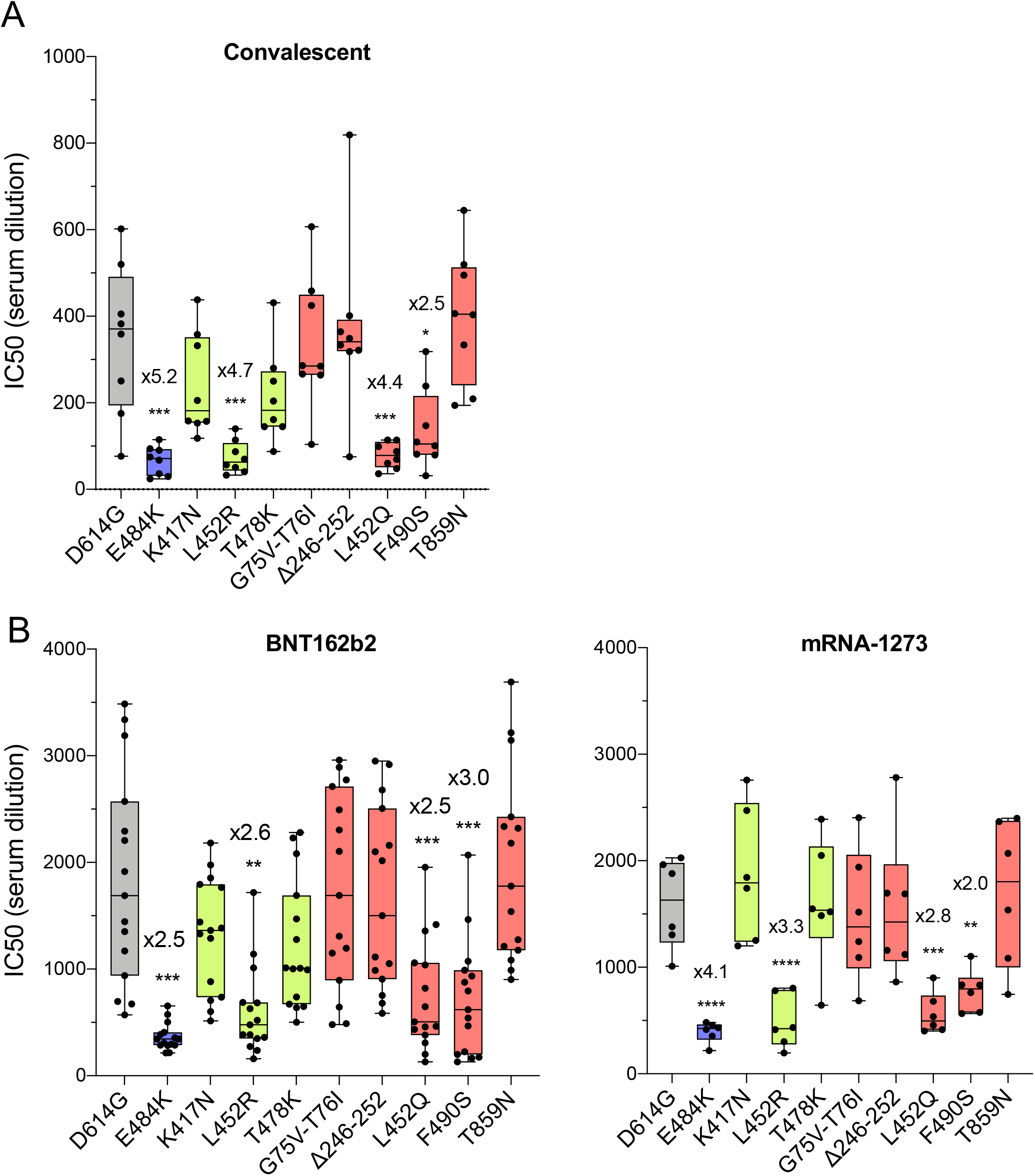
Neutralization titers of spike protein pseudotyped viruses (single point mutations) by convalescent sera, antibodies elicited by BNT162b2, mRNA-1273. (A) Neutralization of variant spike protein (single point mutations) pseudotyped viruses by convalescent serum (n=8). Dots represent the IC_50_ of single donors. (B) Neutralizing titers of serum samples from BNT162b2 vaccinated individuals (n=15). The serum was collected at early time point (7 days after second immunization). Each dot represents the IC_50_ for a single donor. (C) Neutralizing titers of serum samples from mRNA-1273 vaccinated donors (n=6). The serum was collected at early time point (7 days after second immunization). The neutralization IC_50_ from individual donors is shown. Significance is based on the two-sided t-test.

**Supplemental Figure S3.**
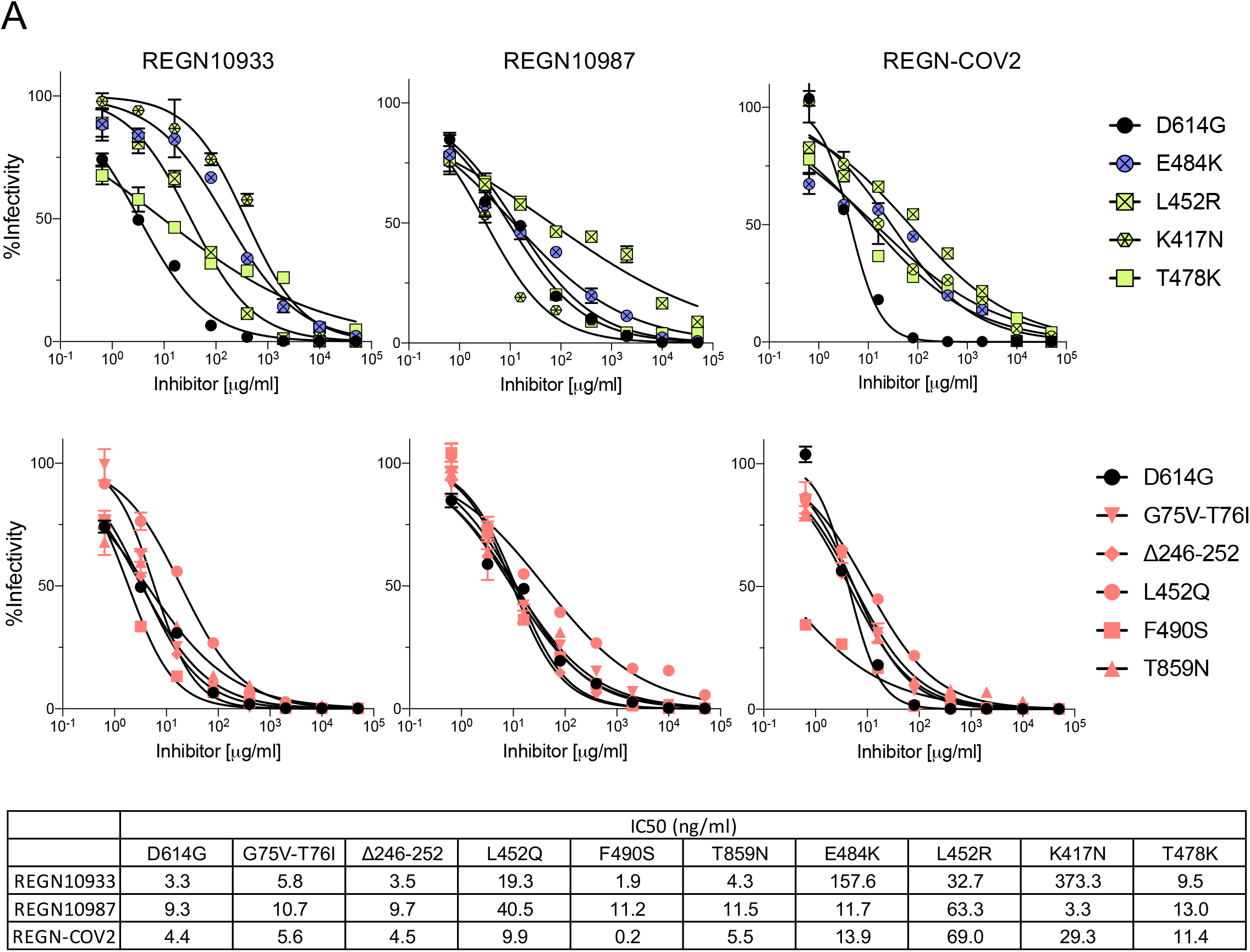
Neutralization titers of spike protein pseudotyped viruses (single point mutations) by monoclonal antibodies. Neutralization of variant spike protein variants (single point mutations) by REGN10933 and REGN10987 monoclonal antibodies. The IC_50_ of REGN10933, REGN10987 and the cocktail is shown in the table.

---

**Table S1.**
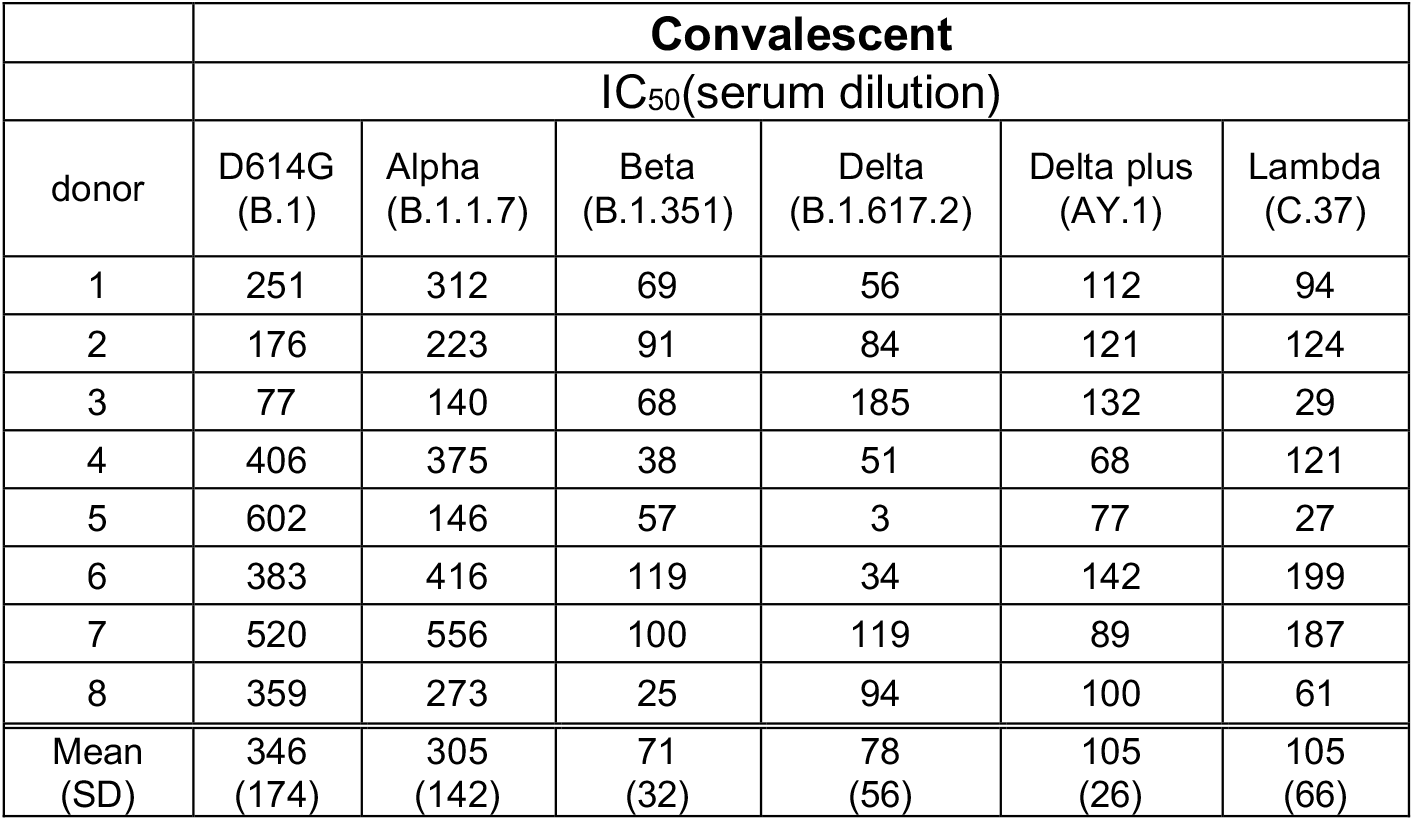
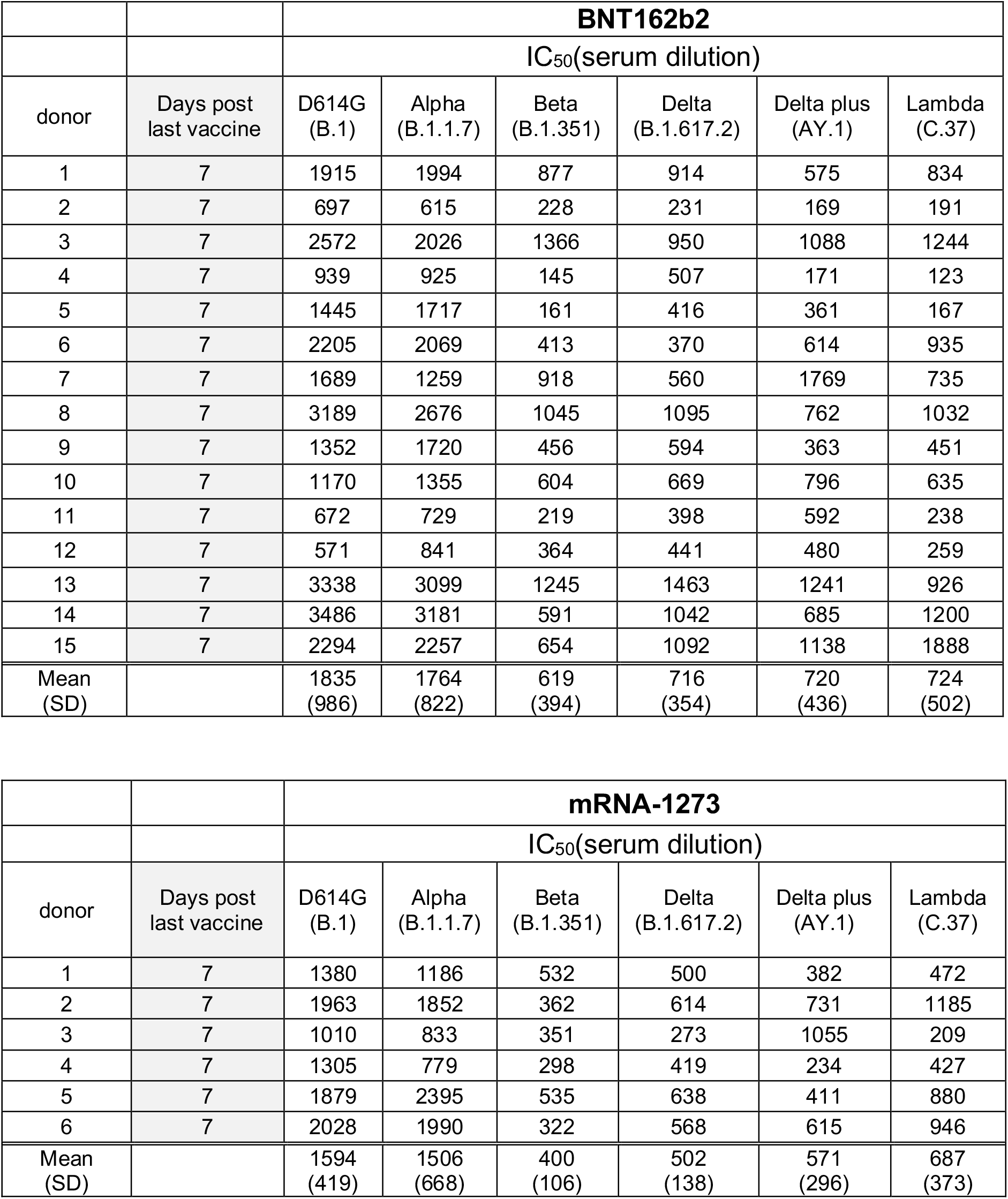
Neutralization of variants by convalescent sera, BNT162b2 and mRNA-1273 elicited antibodies 7 days post-second vaccination.

|  | **Convalescent** |  |  |  |  |  |
| --- | --- | --- | --- | --- | --- | --- |
|  | IC<sub>50</sub>(serum dilution) |  |  |  |  |  |
| donor | D614G<br>(B.1) | Alpha<br>(B.1.1.7) | Beta<br>(B.1.351) | Delta<br>(B.1.617.2) | Delta plus<br>(AY.1) | Lambda<br>(C.37) |
| 1 | 251 | 312 | 69 | 56 | 112 | 94 |
| 2 | 176 | 223 | 91 | 84 | 121 | 124 |
| 3 | 77 | 140 | 68 | 185 | 132 | 29 |
| 4 | 406 | 375 | 38 | 51 | 68 | 121 |
| 5 | 602 | 146 | 57 | 3 | 77 | 27 |
| 6 | 383 | 416 | 119 | 34 | 142 | 199 |
| 7 | 520 | 556 | 100 | 119 | 89 | 187 |
| 8 | 359 | 273 | 25 | 94 | 100 | 61 |
| Mean<br>(SD) | 346<br>(174) | 305<br>(142) | 71<br>(32) | 78<br>(56) | 105<br>(26) | 105<br>(66) |

|  |  | **BNT162b2** |  |  |  |  |  |
| --- | --- | --- | --- | --- | --- | --- | --- |
|  |  | IC<sub>50</sub>(serum dilution) |  |  |  |  |  |
| donor | Days post last vaccine | D614G (B.1) | Alpha (B.1.1.7) | Beta (B.1.351) | Delta (B.1.617.2) | Delta plus (AY.1) | Lambda (C.37) |
| 1 | 7 | 1915 | 1994 | 877 | 914 | 575 | 834 |
| 2 | 7 | 697 | 615 | 228 | 231 | 169 | 191 |
| 3 | 7 | 2572 | 2026 | 1366 | 950 | 1088 | 1244 |
| 4 | 7 | 939 | 925 | 145 | 507 | 171 | 123 |
| 5 | 7 | 1445 | 1717 | 161 | 416 | 361 | 167 |
| 6 | 7 | 2205 | 2069 | 413 | 370 | 614 | 935 |
| 7 | 7 | 1689 | 1259 | 918 | 560 | 1769 | 735 |
| 8 | 7 | 3189 | 2676 | 1045 | 1095 | 762 | 1032 |
| 9 | 7 | 1352 | 1720 | 456 | 594 | 363 | 451 |
| 10 | 7 | 1170 | 1355 | 604 | 669 | 796 | 635 |
| 11 | 7 | 672 | 729 | 219 | 398 | 592 | 238 |
| 12 | 7 | 571 | 841 | 364 | 441 | 480 | 259 |
| 13 | 7 | 3338 | 3099 | 1245 | 1463 | 1241 | 926 |
| 14 | 7 | 3486 | 3181 | 591 | 1042 | 685 | 1200 |
| 15 | 7 | 2294 | 2257 | 654 | 1092 | 1138 | 1888 |
| Mean (SD) |  | 1835 (986) | 1764 (822) | 619 (394) | 716 (354) | 720 (436) | 724 (502) |

|  |  | **mRNA-1273** |  |  |  |  |  |
| --- | --- | --- | --- | --- | --- | --- | --- |
|  |  | IC<sub>50</sub>(serum dilution) |  |  |  |  |  |
| donor | Days post last vaccine | D614G (B.1) | Alpha (B.1.1.7) | Beta (B.1.351) | Delta (B.1.617.2) | Delta plus (AY.1) | Lambda (C.37) |
| 1 | 7 | 1380 | 1186 | 532 | 500 | 382 | 472 |
| 2 | 7 | 1963 | 1852 | 362 | 614 | 731 | 1185 |
| 3 | 7 | 1010 | 833 | 351 | 273 | 1055 | 209 |
| 4 | 7 | 1305 | 779 | 298 | 419 | 234 | 427 |
| 5 | 7 | 1879 | 2395 | 535 | 638 | 411 | 880 |
| 6 | 7 | 2028 | 1990 | 322 | 568 | 615 | 946 |
| Mean (SD) |  | 1594 (419) | 1506 (668) | 400 (106) | 502 (138) | 571 (296) | 687 (373) |

**Table S2.**
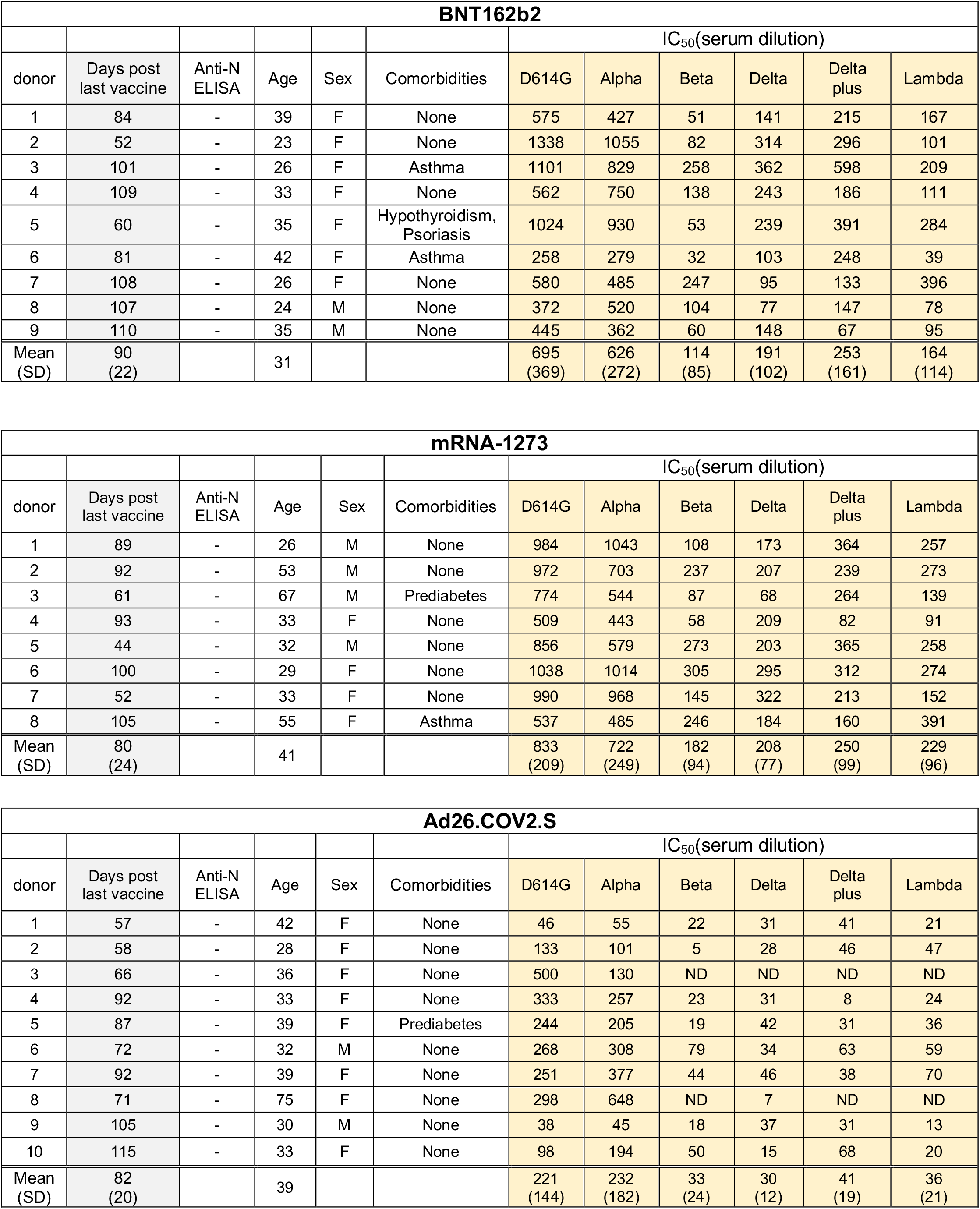
Neutralization of viruses by sera from BNT162b2, mRNA-1273 and Ad26.COV.S vaccinated individuals.

